# Aging Impairs Temporal Integration in Supragranular but Not Thalamorecipient Layers of Primary Auditory Cortex

**DOI:** 10.64898/2026.04.30.722032

**Authors:** Yunru Chen, Patrick O. Kanold

## Abstract

Speech has harmonic features and speech perception is impaired in aging. Imaging of auditory cortex (A1) in aged mice shows reduced selectivity to harmonic sounds and impaired temporal integration of component frequencies especially in layers 2/3 but not layer 4, suggesting that age-related changes in intracortical processing contribute to the hearing deficits. Since complex sounds can be decoded from aging A1, paradigms enhancing cortical processing might improve hearing in aging.

## Main

Age-related hearing impairment reflect both peripheral pathology and central processing deficits^1–3^, which can occur regardless of the peripheral damage. Aging humans have reduced speech perception accuracy^4,5^. Harmonicity is a fundamental feature of speech and animal vocalizations that requires high-fidelity temporal processing of sounds, which is impaired in aging humans^6–8^ likely contributing to their impaired speech perception. The locus of such deficits, however, is unknown. Aging mammals including human, mouse, and rat, show restructured circuits and reduced inhibition in the auditory cortex (ACtx)^9–12^, and these circuit changes could alter the temporal integration of complex sounds^13^. Since a subpopulation of ACtx is exquisitely responsive to harmonic sound emerging at primary ACtx (A1) layer (L) 4^14–16^, we investigated here whether spectro-temporal processing of harmonic stimuli is impaired in aging with a layer-specific manner at L2/3 and L4 as main proxies of intracortical and thalamocortical processing, respectively.

We performed in vivo two-photon imaging in A1 L4 and L2/3 of CBA/CaJ animals that have preserved peripheral hearing^17,18^ (young: 3-6 months; aging: 20-24 months) (Fig. 1a-b, Extended Fig.1a-b) with expression of calcium indicators GCaMP6. We presented pure tone components and two-tone harmonic stacks (fundamental frequency, F_0_: 4-16kHz; Fig. 1c) investigated the response of single neurons and neuronal populations.

**Figure 1.**
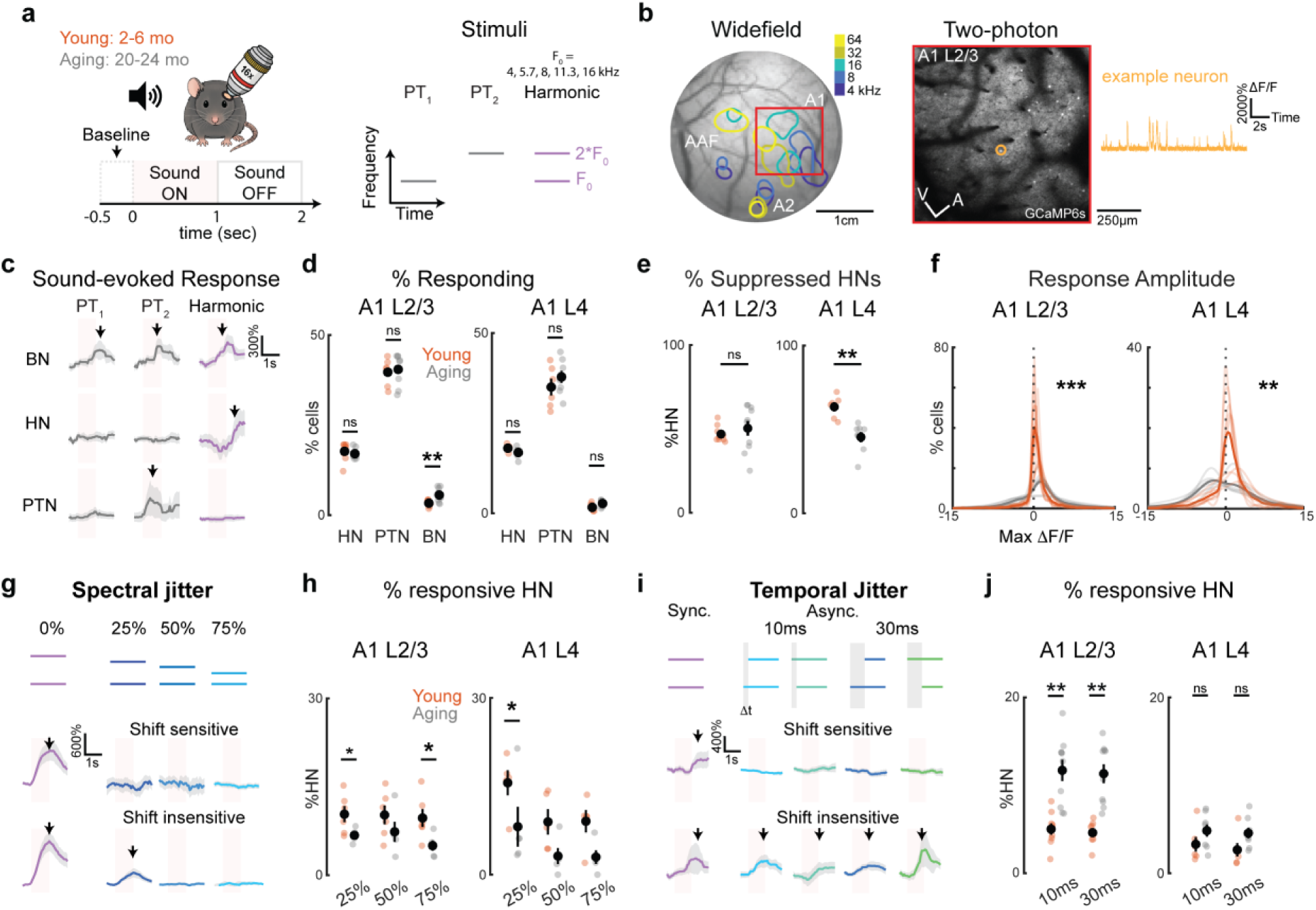
Degraded spectrotemporal tuning of harmonic-sensitive neurons in aging ACtx. **a.** Experiment scheme of in vivo two-photon imaging (left) and the composition of harmonic stacks (right). **b.** An example widefield session showing the most frequency-evoked areas in three subareas. Noting that the evoked response from low to high frequency in A1 follows a tonotopy axis from ventral posterior to dorsal anterior (left). Within the selected A1 area (red rectangle), single neurons (yellow circle) response to harmonic sound (yellow trace) were observed in an example two-photon imaging session (right). **c.** Examples of classified neurons showing distinctly selective response to only harmonic stacks (HN), only pure tones (PTs), or both sounds (BN). **d.** Proportions of classified neurons based on their sound-evoked response showed an significant increase of %BN in aging A1 L2/3. Error bars indicate standard errors (SEM) across animals. The black dot indicates animal average. Individual colored dots indicate one animal in either young (red) or aging (grey) cohort. Total neuron count: YL2/3: 10584 sound-evoked neurons, 9 animals; AL2/3: 19378 neurons, 10 animals; YL4: 3617 neurons, 6 animals; AL4: 7138 neurons, 8 animals. Two sample t-tests were used. YHN, L2/3: 17.70%±0.82%, AHN, L2/3: 17.05%±0.42%, *p*HN, L2/3 = 0.4731. YHN, L4: 18.04%±0.38%, AHN, L4: 16.05%±0.46%, *p*HN, L4 = 0.0830. YPTN, L2/3: 39.67%±1.02%, APTN, L2/3: 40.45%±1.27%, *p*PTN, L2/3 = 0.65. YPTN, L4: 35.00%±2.37%, APTN, L4: 37.82%±1.68%, *p*PTN, L4 = 0.34. YBN, L2/3: 3.32%±0.28%, ABN, L2/3: 5.60%±0.58%, *p*BN, L2/3 = 0.0033. YBN, L4: 1.62%±0.42%, ABN, L4: 2.70%±0.31%, *p*BN, L4 = 0.058. **e.** Proportions of HNs showed a significantly lower %HN with suppressed response to harmonic stacks in A1 L4 of aging animals. Two sample t-tests were used. YL2/3: 46.90%±1.61%, AL2/3: 50.34%±4.39%, *p*L2/3 = 0.49. YL4: 63.49%±2.79%, AL4: 45.41%±3.04%, *p*L4 = 0.0012. **f.** Probability density distribution of the amplitude of ΔF/F at top 95% magnitude (i.e. absolute value) identified in significant sound-evoked response during the response window (1s after sound onset followed by 1s after sound offset). Thicker line indicates the animal-average of individual distributions, thinner line indicates individual distribution. Dotted line indicates ΔF/F at 0. Permutation test comparing area under curve with 10,000 iterations: *p*L2/3 = 0.0003, *p*L4 = 0.0083. **g.** Stimuli design of the imaging sessions probing the spectral integration of harmonic stacks shows that, for each harmonic stack, 25%, 50%, or 75% downward shift was applied the higher frequency in the stack. Example traces show HN with selective response only to the harmonic stacks or with response to both the harmonic stacks and at least one of the shifted stacks. **h.** Quantification of %HN responding to both the original harmonic stack (purple in g) but also any of the spectrally shifted stacks (blue in g) showed significantly decreased %HN responding to the 25% shift and 75% shift in A1 L2/3, and 25% shift in A1 L4. A1 L2/3: Total neuron count: YL2/3: 7120 sound-evoked neurons, 6 animals; AL2/3: 10030 neurons, 5 animals; YL4: 5627 neurons, 5 animals; AL4: 3966 neurons, 5 animals. Two sample t-tests were used. Y25%, L2/3: 10.35%±1.36%, A25%, L2/3: 6.77%±0.51%, *p*25%, L2/3 = 0.048. Y25%, L4: 15.59%±2.13%, A25%, L4: 8.14%±3.37%, *p*25%, L4 = 0.026. Y50%, L2/3: 10.22%±1.59%, A50%, L2/3: 7.36%±1.71%, *p*50%, L2/3 = 0.2515. Y50%, L4: 8.97%±2.17%, A50%, L4: 3.15%±1.36%, *p*50%, L4 = 0.0526. Y75%, L2/3: 9.70%±1.53%, A75%, L2/3: 5.01%±0.87%, *p*75%, L2/3 = 0.033. Y75%, L4: 9.07%±1.90%, A75%, L4: 2.99%±1.17%, *p*75%, L4 = 0.099. **i.** Stimuli design of the imaging sessions probing the temporal integration of harmonic stacks shows that, for each harmonic stack, 10ms or 30ms shift was applied to either the higher or the fundamental frequency. Example traces show an example HN with response to both the synchronized stacks and the asynchronized stacks, as well as an example HN with response only to the synchronized stacks. **j.** Quantification of %HN responding to both the synchronized harmonic stack (purple in **i**) but also the asynchronized stacks (blue and green in **i**) showed significantly increased %HN responding to 10ms- and 30ms-shifted asynchronized stacks in A1 L2/3 but not in A1 L4. Total neuron count: YL2/3: 10584 sound-evoked neurons, 9 animals; AL2/3: 19378 neurons, 10 animals; YL4: 3617 neurons, 6 animals; AL4: 7138 neurons, 8 animals. Two sample t-tests were used. Y10ms, L2/3: 5.01% ± 0.64%, A10ms, L2/3: 11.70% ± 1.22%, *p* 10ms, L2/3 = 2.1052e-4. Y10ms, L4: 3.29%±0.83%, A10ms, L4: 4.83%±0.69%, *p*10ms, L4 = 0.18 Y30ms, L2/3: 4.61%±0.42%, A30ms, L2/3: 11.32%±1.08%, *p*30ms, L2/3 = 3.37e-5. Y30ms, L4: 2.66%±0.77%, A30ms, L4: 4.56%±0.60%, *p*30ms, L4 = 0.071.

We first categorized neurons based on their responsiveness to two distinct sound types: pure tones (PTs) and harmonic (H) stacks, as neurons selective for one of these stimuli (PTN, HN) or responsive to both stimulus types (broadly responsive; BN) at each F_0_. Because individual neurons can exhibit distinct tuning properties across F_0_, we utilized a non-exclusive classification scheme. For instance, a neuron exhibiting PT-exclusive response to F_0_ = 4kHz but responding to both PT and HN to F_0_= 5.3kHz was classified and quantified in both PTN and BN categories. This approach reflects the dynamic capacity of individual neurons to differently encode sound types across a range of F0. By accounting for these overlapping response profiles, we identify subtle shifts in categorical selectivity, such as a neuron gaining PT sensitivity at a specific F_0_ while retaining its harmonic-only response elsewhere, which would have been obscured by a single-label classification. We next examined whether these categorical profiles are altered in the aging ACtx.

The proportion of PTNs and HNs remained stable across young and aging cohorts in both L4 and L2/3 (Fig. 1d). However, the proportion of BNs increased with aging A1 in L2/3 (YBN, L2/3: 3.32%±0.28%, ABN, L2/3: 5.60%±0.58%, *p*BN, L2/3 = 0.0033; Fig. 1d), but not in aging L4 (YBN, L4: 1.62%±0.42%, ABN, L4: 2.70%±0.31%, *p*BN, L4 = 0.058; Fig. 1d). Notably, this expansion of the BN population occurs without a concomitant decrease in PTNs or HNs. This indicates that the aging neurons do not simply switch categories but likely expand their responsive categories at certain frequencies. This localized increase in BNs suggests that L2/3 undergoes a loss of categorical sparsity, consistent with reduced sideband inhibition^19^ that, if preserved, would sharpen spectral tuning and suppress the response at neighboring frequencies. Among the HNs, aging animals have a lower suppressed response to the sound onset in A1 L4 (YL2/3: 46.90%±1.61%, AL2/3: 50.34%±4.39%, *p*L2/3 = 0.49. YL4: 63.49%±2.79%, AL4: 45.41%±3.04%, *p*L4 = 0.0012; Fig. 1e). However, the response profile of HNs across animals showed an expanded range of response amplitude of the aging HNs in both layers (Permutation tests with 10,000 iterations, *p*L2/3 = 0.0003, *p*L4 = 0.0083; Fig. 1f). Altogether, we identified a slightly enlarged population responding to harmonics in aging animals. Fewer HNs in aging A1 L4 showed suppressed onset response, but those with suppressed response had higher amplitude. This suggests a decreased inhibitory regulation of the excitatory response to harmonics in A1 L4, which may, in turn affect the range of response amplitude in both L4 and L2/3.

To validate the criticality of harmonic relationship to the sound-evoked response of HN, we presented a set of two-tone stimuli with down-shifted top frequency in the harmonic stacks by 25%, 50% and 75% (Fig. 1g). In young animals, the fraction of HNs responding to harmonic stacks was reduced to less than 16%, supporting the relatively high sensitivity of HN to the harmonic relationship of two tones in the stack. Aging animals showed decreased proportion of HNs responding to spectral shift in L2/3 and L4 (Fig.1h). This result is consistent with previous studies that show decreased bandwidth^20^ during aging.

In addition to HN’s sensitivity to harmonic relationship, another hallmark of harmonic complexes is the co-occurrence of the different harmonic components. Thus, we tested whether the ability to bind these components together is impaired in aging. We presented harmonic stimuli in which the components’ onsets were temporally shifted by ±10ms or ±30ms relative to F_0_, thereby reducing synchrony. A1 L2/3 of aging animals exhibited a higher proportion of HNs (Y10ms, L2/3: 5.01%±0.64%, A10ms, L2/3: 11.70%±1.22%, *p*10ms, L2/3 = 2.11e-4. Y30ms, L2/3: 4.61%±0.42%, A30ms, L2/3: 11.32%±1.08%, *p*30ms, L2/3 = 3.37e-5. Fig. 1i-j) that remained responsive to temporally shifted asynchronized HM compared to young adults. Such loss of temporal selectivity, however, was not present in A1 L4 (Y10ms, L4: 3.29%±0.83%, A10ms, L4: 4.83%±0.69%, *p*10ms, L4 = 0.18. Y30ms, L4: 2.66%±0.77%, A30ms, L4: 4.56%±0.60%, *p*30ms, L4 = 0.072). Since L4 provides the dominant input to L2/3, the loss of temporal selectivity in L2/3 suggests that central aging affects the intracortical circuitry between L4 and L2/3.

Nonlinear computation is a hallmark of cortical processing, including spectral and temporal integration^16,21–23^. We compared the observed responses to harmonics with a linear reconstruction derived from the same HN’s response to PT components. While young and aging L2/3 neurons exhibited a broad non-normal distribution of R^2^ values for linear reconstructions (Extended Fig.2), aging A1 L2/3 had higher overall R^2^, indicating increased linearity (Permutation tests on Area Under the Differences with 5000 shuffles of animal group, *p* = 0.03; Extended Fig.2). In contrast, no differences were present between ages in L4 (Permutation tests on Area Under the Differences with 5,000 shuffles of animal group, *p* = 0.33; Extended Fig.2). The shift towards increased linearity with age suggests that the aging ACtx loses the specialized nonlinear processing. The similarity at L4 between young and aging animals indicates that the loss of nonlinearity originates within A1 circuits instead of subcortical areas.

**Figure 2.**
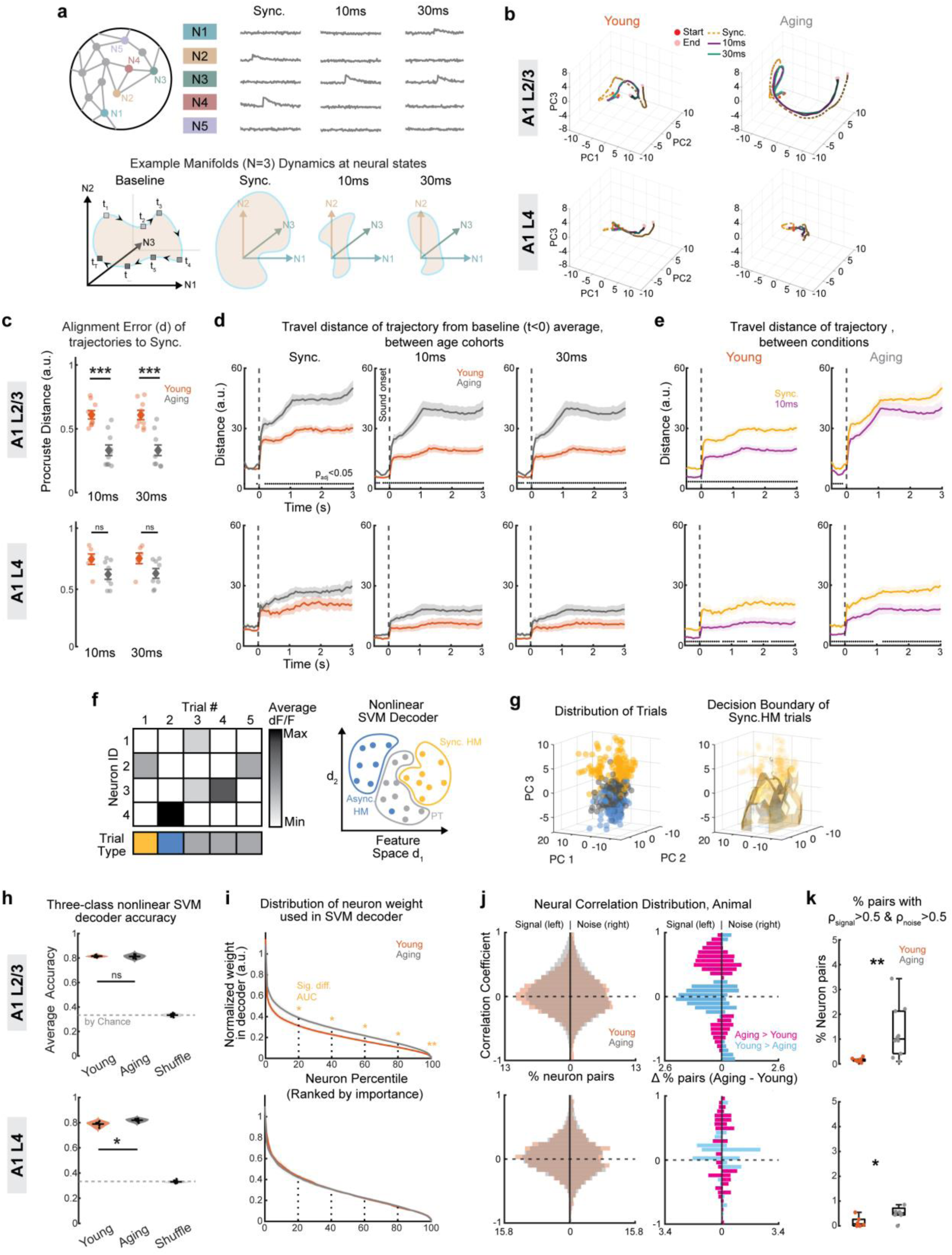
Preserved encoding accuracy of distinct sound types with altered strategy in aging L2/3. **a.** Top: Schematic showing the construction of data matrix including neurons showing or not showing evoked response to harmonics, 10ms-shift, and 30ms-shift stimuli, used to capture the total sound-evoked population. Bottom: example population manifold showing contour at the N-dimension space capturing the activity of N neurons during synchronous harmonics (Sync.), 10ms shift (10ms) and 30ms shift (30ms) conditions. **b.** Animal-average population trajectories in neural state space in A1 L2/3 (top) and L4 (bottom) for young (left) and aging (right) animals, shown from start (saturated red dot, sound onset) to end (light red dot, 2s after sound onset), including Sync. (yellow), 10ms-shift (purple), and 30ms-shift (green) conditions. Shading indicates light (early), mid, and dark (late) time after sound onset. **c.** Quantification of the alignment error (Procrustes distance) between trajectories of 10ms-shift and 30ms-shift conditions relative to the Sync. trajectory in young and aging cohorts, in A1 L2/3 (top) and L4 (bottom). Two sample t-tests were used. Y10ms, L2/3: 0.61±0.03, A10ms, L2/3: 0.33±0.04, *p*10ms, L2/3 = 7.13e-5. Y30ms, L2/3: 0.61±0.03, A30ms, L2/3: 0.33±0.04, *p*30ms, L2/3 = 7.14e-4. Y10ms, L4: 0.75±0.04, A10ms, L4: 0.62±0.04, *p*10ms, L4 = 0.06. Y30ms, L4: 0.75±0.04, A30ms, L4 0.63±0.04, *p*30ms, L4 = 0.06. * *p*00.05, ** *p*00.01, *** *p*00.001. **d.** Travel distance of neural trajectories from the mean of baseline (i.e. average ΔF/F within 0.5s before sound onset) across age cohorts for Sync. (left), 10ms-shift (middle), and 30ms-shift (right) conditions in A1 L2/3 (top) and L4 (bottom). Significantly higher errors were observed in A1 L2/3 aligning 10ms-shift or 30ms-shift trajectories to Sync. Shaded areas indicate SEM. Solid lines indicate animal averages of young (red) or aging (gray) cohorts. Wilcoxon rank sum tests were used to compare young and aging cohorts at each time point. Black dot indicates *p* 0 0.05 after Benjamini-Hochberg correction. **e.** Similar to **d** but travel distance of neural trajectories compared between conditions (Sync, yellow; 10ms-shift, purple) for young (left) and aging (right) animals in A1 L2/3 (top) and L4 (bottom). Wilcoxon rank sum tests were used to compare young and aging cohorts at each time point. Black dot indicates *p* 0 0.05 after Benjamini-Hochberg correction. **f.** Left: example input data matrix to a nonlinear SVM decoder composed of average ΔF/F of each neuron within the response window (i.e. 1s after sound onset and 1s after sound offset) for each trial of every unique sound. Right: example 2-dimension feature space showing classification of three trial types being PT (gray), synchronous harmonics (yellow) and asynchronous harmonics (blue). The color of the dot indicates the ground truth identity of the trial. Colored boundaries indicate the classification result from the nonlinear SVM decoder. **g.** Left: example 3-dimension PCA space showing the distribution of true trial types. Right: nonlinear hyperplane (yellow) as the decision boundary of synchronous harmonics trials separates the synchronous harmonics trials from other trial types. **h.** Three-class nonlinear SVM decoder accuracy in A1 L2/3 (top) and L4 (bottom) for young, aging and shuffle groups. Mann-Whitney U tests were used followed by Benjamin-Hochberg correction. * Indicates *p* 0 0.05 after correction. YL2/3: 80.91% ± 0.35%, AL2/3: 82.23% ±0.60%, ShuffleL2/3: 33.29%±0.18%, *p*Y vs A, L2/3 = 0.1160, *p*Shuffle vs A, L2/3 = 1.44e-5, *p*Y vs Shuffle, L2/3 = 2.88e-5. YL4: 79.53% ±0.91%, AL4: 81.98% ±0.42%, ShuffleL4: 33.26%±0.28%, *p*Y vs A, L4 = 0.0026, *p*Shuffle vs A, L4 = 1.51e-4, *p*Y vs Shuffle, L4 = 2.88e-5. **i.** Distribution of normalized neuron weights used in the full linear SVM decoder, showing that aging cohort had significantly higher area under curves at the top 10%, 20% and 30% important neurons used by the linear SVM decoder. Shaded Area indicates SEM of he weight distribution curve. Dashed vertical line at N indicates comparisons of area under curve from top 0% to top N% between young and aging cohorts. Two sample t-tests were used. * *p*00.05. **j.** Left: Animal-average histograms of signal and noise correlation coefficients across neuron pairs in A1 L2/3 (top) and L4 (bottom) for young (red) and aging (gray) animals. Right: Differences of the animal-average histograms of young and aging cohorts for signal correlations (top) and noise correlations (bottom). Magenta, aging > young. Blue, young 0 aging. **k.** Quantification of the proportion of most correlated neuron pairs (with both signal and noise correlations larger than 0.5) among all pairs of sound-evoked neurons showing a significantly higher proportion of most correlated pairs in aging animals in both A1 L2/3 and L4. Mann-Whiteney U tests were used. YL2/3: 0.6% ± 0.2%, AL2/3: 1.7% ± 0.5%, *p*L2/3 = 0.012. YL4: 0.2%±0.1%, AL4: 1.2%±0.8%, *p*L4 = 0.041.

So far, we have shown the degradation of spectrotemporal integration and encoding of HM at the single-neuron level. However, the brain’s ability to recognize and categorize complex sounds relies on the collective activity of neuronal ensembles, and population-level encoding accounts for the high-dimensional interactions and redundancies^24,25^. By employing neural manifold alignment approach based on Procrustes analysis (Fig, 2a), we characterized the population-level dynamics responding to synchronized harmonics and temporally shifted harmonics, respectively, for L4 and L2/3 in each animal (Fig. 2b). To visualize these complex population-level dynamics, we projected the high-dimensional ensemble activity into a lower-dimensional manifold using Principal Component Analysis (PCA). By capturing the dominant modes of variance, this allowed us to represent the temporal evolution of the neural response as trajectories within a common PC space. In young animals the trajectories in L2/3 ensembles in PC space diverged for synchronized harmonics and 10ms- or 30ms-shifted stimuli, indicating that neural population activity differed for these stimuli along their duration. In contrast, in aging L2/3 the trajectories occurred over a larger range in PC space and appeared more similar for all 3 sound conditions. To compare the neural representation of jittered stimuli (*H*_10*ms*_, *H*_30*ms*_) to the synchronous template (*H*_*sync*_), we used Procrustes superimposition. For each F0, the *H*_*sync*_trajectory served as the fixed reference manifold. The trajectories for the *H*_10*ms*_ and *H*_30*ms*_ conditions were then translated, rotated, and scaled to align with the corresponding *H*_*sync*_ template. The degree of manifold divergence was quantified using the Procrustes distance *d*. A larger *d* indicates that temporal jittering induced a more profound reorganization of the population dynamics that could not be reconciled through linear transformations. Aging A1 L2/3 showed decreased alignment error between trajectories of sound types (Y_10ms, L2/3_: 0.61±0.03, A_10ms, L2/3_: 0.33±0.04, *p*_10ms, L2/3_ = 7.13e-5. Y_30ms, L2/3_: 0.61±0.03, A_30ms, L2/3_: 0.33±0.04, *p*_30ms, L2/3_ = 7.14e-4; Fig.2c), while A1 L4 remained more stable (Y_10ms, L4_: 0.75±0.04, A_10ms, L4_: 0.62±0.04, *p*_10ms, L4_ = 0.06. Y_30ms, L4_: 0.75±0.04, A_30ms, L4_: 0.63±0.04, *p*_30ms, L4_ = 0.06; Fig.2c). This indicates that the population response becomes less synchronization-specific in the aging L2/3, potentially contributing to age-related deficits in the detection and categorization of sounds.

With the use of kinematic metrics on each time step of the temporal trajectory, we found that aging A1 L2/3, but not L4, showed an increased distance from baseline across all stimulus conditions (black dot indicates *p*00.05; Fig 2d). At a closer look comparing travel distance at synchronous and 10ms conditions, young animals showed a significantly larger distance traveled during synchronous condition than the 10ms-shift condition, which is not observed in aging animals (black dot indicates *p*00.05; Fig 2e). The overall increase of travel distance in aging L2/3 trajectories indicate the neuron population is more broadly activated regardless of stimulus type. Specifically, while young animals selectively travel further for synchronous condition than 10ms-shift, the travel distances do not differ between the two conditions in aging animals. This result suggests a reduced stimulus specificity in aging animals as the neural state space no longer differentiates synchronous from temporally shifted stimuli as effectively. Notably, this is only observed in L2/3 but not L4, marking the emergence of degraded differentiation at the cortical stage. Additionally, aging cohorts also showed increased peak velocity of the trajectory (black dot indicates *p*00.05; Extended Fig. 3a) across all conditions. Both young and aging animals showed increased velocity during synchronous condition than 10ms-shift condition in L2/3 and L4 (Extended Fig. 3b). The increased velocity indicates a faster rate of change in the state-space by the neuron population over time and suggests the collective firing pattern of aging A1 L2/3 is traveling more rapidly from one state to another as a sign of decreased stability between neural states. We speculated that this manifold collapse reflects a fundamental shift in the population coding ability that 1) may simply be the outcome of degraded temporal acuity, or 2) serves as active compensation for reduced single-neuron selectivity to temporal features in the harmonic sounds. Above, we revealed that, within the PCA subspace constructed by population activities to synchronous and asynchronous harmonics, aging populations show expanded trajectories but with greater overlap between conditions (Fig. 2a-e). Here we applied PCA to population activities across the stimulus set (i.e., PTs, synchronous and asynchronous harmonics) and acquired activity patterns, or PCs observed across stimuli (Extended Fig.4). Aging L2/3 require more PCs to explain 95% of variance, indicating that representations across the full tonal and temporal stimulus space are distributed across more dimensions. The difference is more pronounced in the early response window (0.3s after onset) compared to a longer response window (1s after onset), suggesting that the degradation of low-dimensional population structure in aging is most evident during the initial stimulus-driven response rather than sustained activity. Together, these findings suggest a larger but less organized population code in aging A1 L2/3.

To test if the change in ensemble activity was still able to accurately represent the stimuli, we employed Support Vector Machine (SVM) decoders to classify trial types to either “*H*_*sync*_”, “*H*_*async*_” or “PTs” (Fig.2f-g). We trained a three-class nonlinear SVM and despite the observed manifold collapse, the decoder achieved similar average accuracies across cohorts in both layers (Y_L2/3_: 80.91% ±0.35%, A_L2/3_: 82.23% ±0.60%, Shuffle_L2/3_: 33.29%±0.18%; Y_L4_: 79.53% ±0.91%, A_L4_: 81.98% ±0.42%, Shuffle_L4_: 33.26%±0.28%; Fig. 2h), suggesting a robust compensatory encoding mechanism in the aging L2/3. To investigate the nature of this compensation, we extracted neuron weights from a linear three-class SVM, in which aging cohorts showed slightly higher accuracies in both layers (Y_L2/3_: 80.93% ±1.12%, A_L2/3_: 87.65% ±1.07%, *p*_L2/3_ = 1.30e-4. Y_L4_: 73.78% ±3.56%, A_L4_: 80.21% ±1.93%, *p*_L4_ = 0.15; Extended Fig. 5), which may result from increased linearity in neuron computation (Extended Fig. 2). In ACtx L2/3 of young animals, classification was dominated by a sparse subset, or the “vital few”, with significantly higher normalized weights emerging at the top 20% neurons (Fig. 2i). In contrast, the aging ACtx L2/3 distributes weights more uniformly across a larger neuron population, indicating a shift toward a more redundant and distributed population representation. Converse to L2/3, A1 L4 remain similar across cohorts (Fig. 2i). The redundant coding is also seen in the increased proportion of most-correlated neuron pairs, prominently in aging L2/3 and in L4. A higher percentage of neuron pairs showed high signal correlation, indicating high similarity in their calcium response to harmonic stimuli, as well as high noise correlation, indicating greater functional connectivity (Fig. 2j-k).

Here, we unveiled A1 L2/3-specific age-related changes on the spectrotemporal integration of harmonic stacks, which are fundamental features of human speech and animal vocalization. These changes are not present in L4, indicating a specific change in intracortical circuits with aging that impairs sound processing^26,27^. Our SVM analysis shows that, in aging, despite deficits in the encoding of harmonic sound features by individual L4 cells, L4 population activity can still encode these features, though the encoding scheme has changed. These results are consistent with increased correlated activity between neurons’ tone-driven activities in aging within L2/3^20^. Current hearing aids focus primarily on peripheral amplification, yet our data suggests that simply increasing gain is insufficient if the central circuits to L2/3 are altered. Given the capacity of cortical circuits to be plastic^28,29^, neuroplastic interventions in combination with next-generation neuroprosthetics may move beyond simple amplification toward compensating or preventing the central deficits. Indeed, musical training seems protective in humans^30,31^ and targeted training paradigms have shown effects in aging mice^32,33^.

## Methods

### Animals

All experimental procedures were approved by the University of Maryland or the Johns Hopkins University Institutional Animal Care and Use Committee. We used the total of *N* = 19 mice (9 young (5F, 4M, 3-6 months old), 10 aging (5F, 5M, 20-24 months old) on a CBA/CaJ background (JAX strain #: 000654) which show similar levels of age-related peripheral hearing loss to humans allowing for the isolated study of central auditory system dysfunction^34–37^ with minimal peripheral hearing loss as compared to the common C57BL/6J strain^34^. Mice in the young cohort are F1 of CBA/CaJ (JAX strain #: 000654) x B6;Thy1-GCaMP6s (JAX strain #: 024275) and mice in the aging cohort are a mixture of the F1 of CBA/CaJ x B6;Thy1GCaMP6s with transgenic expression of calcium indicator (N=3) and with expression by virus injection of AAV.CaMKII.GCaMP6s.WPRE.SV40 (N=7). All analyses in this study are conducted after validating no differences between the transgenic group and the injected group within the aging cohort. Mice with injections were imaged at least one month after the injection. All mice were housed in a 12 h reverse light/dark cycle room, and ambient noise in the room was minimized. Imaging experiments were only performed during the dark cycle.

### Craniotomy and window implantation

Craniotomy and window implantation were performed as described previously^38,39^. About 1 hour before surgery, we injected dexamethasone (1 mg/kg, VetOne) subcutaneously (s.c.) to minimize brain edema. Anesthesia was induced with 4 % isoflurane (Fluriso, VetOne) with a calibrated vaporizer (Matrix VIP 3000) and then maintained at 1.5% - 2% for young animals and 1% - 2% for aging animals. The animal was kept on a heating pad, and its body temperature was monitored using a rectal probe throughout the surgery and maintained at around 36.5°C (Harvard Apparatus Homeothermic System). Hair on the head was first snipped then removed with a hair removal product (Nair). Betadine and 70 % ethanol were applied 3 rounds to the exposed skin. Skin covering the skull was removed to expose the skull of both left and right hemispheres. After removing the skin, connective tissue and the underlying muscles were scraped to the temporal sides under the remaining skin. A unilateral craniotomy was performed to expose an about 3.5 mm diameter region over the left ACtx. Two circular glass coverslips (one 3 mm and one 4 mm in diameter) were affixed with a transparent silicone elastomer (Kwik-sil, World Precision Instruments). A custom 3D-printed stainless steel headpost was attached to the skull above the right hemisphere, and the rest of the exposed skull area and the edge of the cranial window were covered using dental acrylic (C&B Metabond). Meloxicam (5 mg/kg) and cefazoline (300 mg/kg) were injected post-operatively. Animals had at least 14 days before any imaging was performed.

### Virus Injection

After the craniotomy, mice were injected with 300 nL of AAV.CamKII.GCaMP6s.WPRE.SV40 (Addgene viral prep #107790-AAV9) virus at each site and a total of 3 sites near tentative A1 to ensure the full coverage of A1. The injection was done at 400 µm below the pial to deliver GCaMP6s to excitatory neurons^40^. For each injection, a custom pulled (Sutter Instrument Co. P-2000), glass pipette was positioned so that the tip was first staying 450 µm below the pial surface for two minutes then retrieved to 400 µm for actual injections. Virus was then dispensed (Drummond Nanoject III) at a rate of 180nL/minute until the desired amount was dispensed and the pipette remained in place for five additional minutes before being retracted. Once the injection was completed, cranial window was implanted as described above.

### Widefield imaging

Widefield imaging was performed as previously described^41^. Animals were restrained in a 3D-printed plastic tube and head-fixed using a custom designed headpost holder. In vivo widefield imaging was performed by sending 470 nm LED light (Thorlabs, #M470L3) over ACtx using a camera (PCO edge 4.2) to capture 330 x 330 pixel sized images covering the 3 mm diameter cranial window at 30 Hz frame rate. A series of 100 ms pure tones between 4 – 64 kHz in 40, 55, 70dB SPL were presented at a random order to identify regions of ACtx, tracing auditory responsive green fluorescence change. Image processing was performed as previously described^41^. Briefly, on a downsampled image sets by a factor of 3, we applied an autoencoder neural network to run an automatic image segmentation to group pixels with strong temporal correlations as a single component (Regions of Interests, ROIs) by applying a dimensionality reduction ^42^and identified 50 ROIs. For each ROI, images of 15 frames from the sound onset (F_t_) were baseline corrected with the average value of 5 frames before each onset to trace fluorescence changes to different frequency-sound level pairs (F_norm_ = (F_t_ -F_0_) / F_0_). We defined subregions of ACtx from a tonotopic gradient of tone responses for each subregion for Thy1 cells^41^ and similarly for those with virus expressing of GCaMP6s. Here we focus on A1. For A1, a low-to-high frequency gradient (tonotopy) was organized following the caudal to dorsomedial direction.

### Stimuli design

Stimuli used in this study include two major types: pure tones (PTs) and harmonic stacks (H). Two sets of experiments were carried out at each cortical region to investigate the spectral integration and temporal integration, respectively. During the probing of temporal integration, HM stimuli were composed of two harmonic frequencies with the fundamental frequencies being 4kHz, 5.7kHz, 8kHz, 11.3kHz, and 16kHz. PT stimuli include all of the component frequencies presented in any HMs. During the probing of spectral integration, HM stimuli were composed similarly to the temporal integration, except the fundamental frequencies include 4kHz, 5.7kHz, 8kHz, 11.3kHz, 16kHz, 22.6kHz and 32kHz. Spectrally shifted HMs include the same fundamental frequencies with higher frequency down-shifted by 25%, 50%, or 75%. For example, a HM with fundamental frequency of 4kHz would be probed and compared with three spectrally shifted stacks with top frequencies being 7kHz (25%), 6kHz (50%), or 5kHz (25%). For all stimuli, each frequency was generated at 60dB ± 3dB and the two-tone stacks at 63dB ± 3dB.

### In vivo two-photon imaging

At least one day after performing widefield imaging, animals were awake and head-fixed on a custom-made stage. A free field speaker (TDT ED1) was fixed on a holder 10 cm from the animal’s right ear at 45°. All sound stimuli were delivered through TDT RX6 multiprocessor. In-vivo 2p Ca^2+^ imaging was performed over ACtx using Bruker 2P Plus microscope tilted at 48–55 degrees with a resonant scanning mode (Bruker Ultima 2Pplus; 940 nm excitation wavelength). We imaged a field of view (about 1000 μm ×1000 μm) using a Nikon LWD 16x Objective (0.80 NA, 3.0 WD), femtosecond laser (Coherent Discovery at 940 nm), and PrairieView software (version 5.6) at 15 Hz frame rate. For A1 L2/3, we imaged at a depth between 150 and 200 μm below the pia surface. For A1 L4, we imaged at a depth of 360-400 μm below the pia surface. One imaging session lasted 0.5-1 hr. Each animal underwent two imaging sessions at L2/3 and L4, respectively. Each condition includes no more than one imaging session per animal.

### Preprocessing of two-photon imaging data

Motion correction, cell detection, and cell fluorescence traces, including neuropil fluorescence, were extracted from raw imaging data using suite2p software^43^. Any further analyses were done using Matlab (R2025b). Among the detected cells, we selected cells that show sound-evoked responses, following our previously used method^41,44^. Neuropil correction was computed first following the equation: F_cell_corrected_ = F_cell_ – (0.9 * F_neuropil_). Then, we computed ΔF/F by subtracting the average baseline period before the sound onset (500 ms) from F_cell_corrected_, and then dividing by the baseline. To select cells that are responsive with facilitated response to the sound onset, we calculated the 95 % confidence interval (CI) of baseline (500 ms before the sound onset) and sound-evoked activity (1 sec after the sound onset). Cells were regarded as sound-responsive with facilitative response when the lower bound of CI during the response window was above the upper bound of CI during baseline. The suppressed response was identified when the upper bound of CI during the response window was below the lower bound of CI during baseline. To identify cells responsive to the sound offset, we set the baseline as 500ms before the sound offset and only considered facilitated response to the offset when the same criteria were met.

### Categorization of neurons based on sound-evoked response

After identifying sound-evoked response in neurons, we then characterize neurons into three categories: HN (harmonic-selective, no response to the PT components of the played harmonic complex), PTN (PT-selective, no response to the harmonic stimulus composed of the corresponding pure tones), and/or BN (both responding). Given that individual neurons can exhibit heterogeneous tuning properties depending on the F_0_, we implemented a non-exclusive classification scheme. Each neuron was independently evaluated at every F_0_ within its responsive range, allowing a single unit to be represented in different functional categories across the spectral domain. The quantification of proportion of categories was done by dividing the total number of neurons in each category by the total number of sound-evoked neurons.

### Support Vector Machine Decoders

For each mouse, ΔF/F transients from all neurons with sound-evoked response to any sound stimulus were extracted within the response window (1s after onset and 1s after offset). Trial-by-trial mean responses were computed for each neuron to construct a feature matrix *X* ∈ ℝ^*T* × *N*^, where *T* is the number of trials and *N* is the number of neurons. To ensure that neurons with higher firing rates did not disproportionately influence the decoder, features were z-score standardized across trials. To prevent the decoder from developing a bias toward the most frequent stimulus category (“Others”), we performed class balancing using the Synthetic Minority Over-sampling Technique (SMOTE)^45^. Synthetic examples were generated for the minority class (Synchronized harmonic) to match the trial count of the majority classes (10ms-shifted harmonic and 30ms-shifted harmonic). Decoding was performed using an Error-Correcting Output Codes (ECOC) model with Matlab function “fitcecoc”, which utilized three-classes SVM learners with a Gaussian (RBF) kernel with Matlab function “templateSVM”. The kernel scale was automatically heuristic-optimized, and a box constraint of 10 was applied to regulate the margin hardness. Model performance was evaluated using 6-fold cross-validation. In each fold, the data was partitioned into training and testing sets, and the mean decoding accuracy was calculated as:

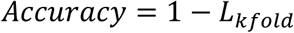

where *L*_*kfold*_ represents the classification loss across all folds.

To establish the statistical significance of the decoding results, we compared the observed accuracy against a null distribution generated through permutation testing. For each mouse, the stimulus labels were randomly shuffled (n=5 iterations), and the decoding pipeline was repeated. The true decoding accuracy was considered significant only if it substantially exceeded the mean accuracy of the shuffled control, which represents the chance-level performance for the balanced dataset (33.3% for a three-class problem).

To identify the individual neurons contributing most significantly to categorical discrimination, we analyzed the weight vectors (coefficients) of the SVM decoder. For each mouse, a linear three-classes SVM decoder as well as a“full” non-partitioned ECOC model were trained using a linear SVM template (Matlab function “templateLinear” with Ridge (L2) regularization. For the full models, each three-classes learner within the ECOC model, we extracted the coefficient vector *β*, where the magnitude |*β*_*i*_ | represents the contribution of neuron *i* to that specific classification boundary. A global “Importance Index” was calculated for each neuron by averaging the absolute weights across all internal binary learners:

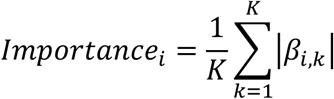

where *K* is the number of binary classifiers. This metric provides a normalized measure of how much a single neuron’s activity influences the overall three-class categorical decision.

To compare the concentration of categorical information between young and aging cohorts, importance weights were normalized to the maximum weight within each mouse 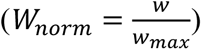 Neurons were then ranked by their importance, and we generated group-averaged importance curves by interpolating the ranked data onto a common percentile-based x-axis (0% to 100% of the population).

### Signal and Noise Correlation

Signal correlations are the synchronized activity between neurons for each stimulus type. We computed the cross-correlation of the average ΔF/F traces across trials, for each stimulus type, between two neurons using the Matlab function “corr”. Noise correlations are the similarity of the trial-to-trial fluctuation between neurons per stimulus type. To calculate noise correlations, we applied the same correlation function on a vector of mean-subtracted amplitudes during the sound presentation per trial level between two neurons for each stimulus type, using the Matlab function “corr”. We compared the distribution of signal/noise correlation coefficients for individual synchronized harmonic stacks.

### Principal Component Analysis

We performed Principal Component Analysis (PCA) on the sound-evoked neuron activity matrices to further look at %variance explained by a range of numbers of principal components. For each animal, a population response matrix *X* ∈ ℝ^*N* × *S*^ was constructed, where *N* represents the number of sound-evoked neurons and *S* represents the total number of unique sound stimuli. The value of each entry (*i*, *j*) was defined as the mean calcium transient (ΔF/F) for neuron *i* during the response window (0.33s or 1s after sound onset). These durations were chosen to capture the response to represent different response window of the played sound. PCA was applied to the population matrices by using the Matlab function “pca”. We calculated the percentage of variance explained by each individual PC and the cumulative variance explained as a function of the number of PC included. To compare population dimensionality between young and aging cohorts, we extracted the average as well as the 95% CI of cumulative variance curves across mice for young and aging cohorts, respectively. For 10, 20, and 30 principal components, respectively, we compared the cumulative variance explained between the two cohorts using a Wilcoxon rank-sum test.

### Statistical Analysis

Statistical evaluations were performed using MATLAB (R2025b, MathWorks). Data distributions were first assessed for normality using the Lilliefors test to determine the appropriateness of parametric versus non-parametric models. For simple comparisons of scalar metrics between Young and Aging groups (e.g., mean decoding accuracy per animal or animal-wide mean importance weights), two-tailed independent samples t-tests were utilized. In cases where the assumption of normality was violated, the Mann-Whitney U test was used as a non-parametric alternative. To compare distribution curves, such as the normalized importance weight profiles, we employed Linear Mixed-Effects (LME) models. In these models, “Age Group” and “Percentile Rank” were treated as fixed effects, while “Mouse ID” was included as a random effect to control for inter-subject variability. To identify specific differences between the Young and Aging curves, we utilized a non-parametric permutation test. The group labels (Young vs. Aging) were randomly shuffled for 5,000-10,000 times to generate a null distribution of the difference between the curves. The observed point-by-point difference was then compared against this null distribution. Multiple comparisons across the curve were corrected using (e.g., False Discovery Rate (FDR) or a cluster-based permutation approach) to maintain a global significance level of *⍺* = 0.05. For all tests, a corrected p-value of less than 0.05 was considered statistically significant.

### Population Trajectory Analysis

For each animal, neural activity was organized into population response matrices *R* ∈ ℝ^*T* × *N*^, where *T* represents the number of time frames and *N* the number of all sound-evoked neurons. To ensure equal weighting of all units, the trial-averaged activity of each neuron was z-score standardized. We performed dimensionality reduction using Principal Component Analysis to extract the top 20 latent dimensions which captured the dominant temporal dynamics of the population response. To compare the neural representation of jittered stimuli (*H*_10*ms*_, *H*_30*ms*_) to the synchronous template (*H*_*sync*_), we used Procrustes superimposition by using the Matlab function “procrustes”. For each F_0_, the *H*_*sync*_ trajectory served as the fixed reference manifold. The trajectories for the *H*_10*ms*_ and *H*_30*ms*_ conditions were then translated, rotated and scaled to align with the corresponding *HM*_*sync*_ template. The degree of manifold divergence was quantified using the Procrustes distance, which represents the residual sum-of-squares between the aligned manifolds. A larger *d* indicates that temporal jittering induced a more aggressive reorganization of the population dynamics that could not be reconciled through linear transformations. The distance from baseline was the Euclidean distance between the neural state at time *t* and the mean baseline coordinates before sound onset (Z_base_), which captures the overall magnitude of the stimulus-evoked excursion in state-space. The velocity (*v*), or the instantaneous rate of change of the neural trajectory was calculated as 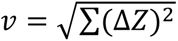, representing the speed of population state transitions. The velocity profiles were then smoothed using a Gaussian kernel (*σ* = 5). For all manifold metrics (Procruste distance, distance from baseline, and velocity) were first averaged across all tested fundamental frequencies within each animal. Group-level comparisons between young and aging cohorts were performed using two-tailed independent samples t-tests. For time-resolved kinematic metrics (distance from baseline and velocity curves), statistical significance at each time point was analyzed by t-tests, followed by Benjamini-Hochberg False Discovery Rate correction to account for multiple comparisons across frames.

## Acknowledgements

YC and POK designed research. YC performed imaging experiments and analysis. YC and POK drafted and edited the manuscript. Supported by NIH R01DC017785 and RF1AG078378. We would like to thank Dr. Minzi Chang, Dr. Didhiti Mukherjee, Dr. Nicholas Michelson for their thoughtful comments on this manuscript.

---

**Extended Figure 1:**
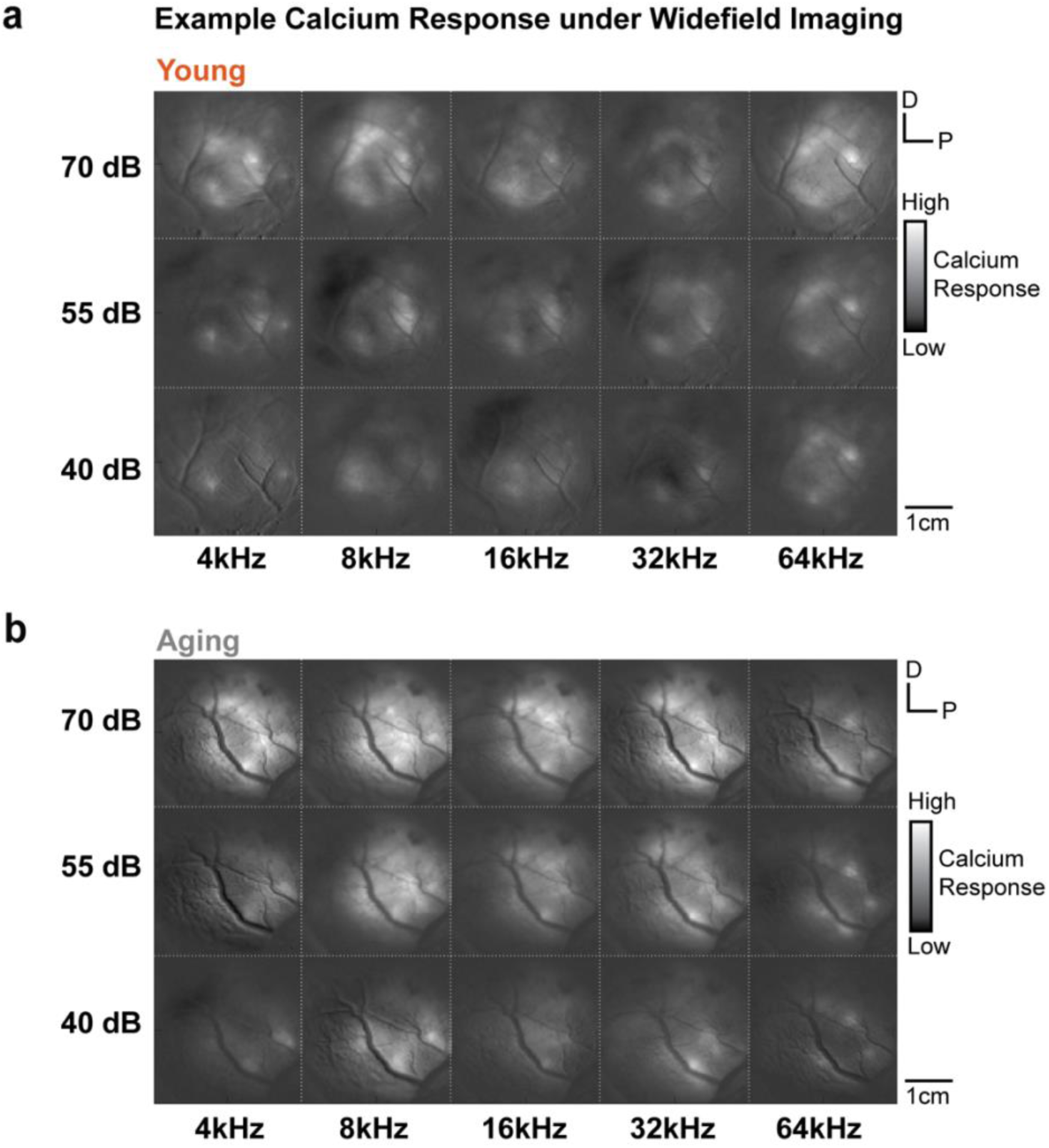
Sound-evoked response of an example young animal and an example aging animal. **a.** Widefield calcium imaging maps showing sound-evoked cortical responses in an example young animal across five frequencies (4, 8, 16, 32, 64 kHz) and three sound levels (40, 55, 70 dB SPL). Color scale indicates calcium response magnitude (high to low). D, dorsal; P, posterior. Scale bar, 1cm. **b.** Same as **a**, but for an example aging animal. Animals from the aging cohort that showed no visible cortical activation at 40dB SPL were excluded from further analysis.

**Extended Figure 2:**
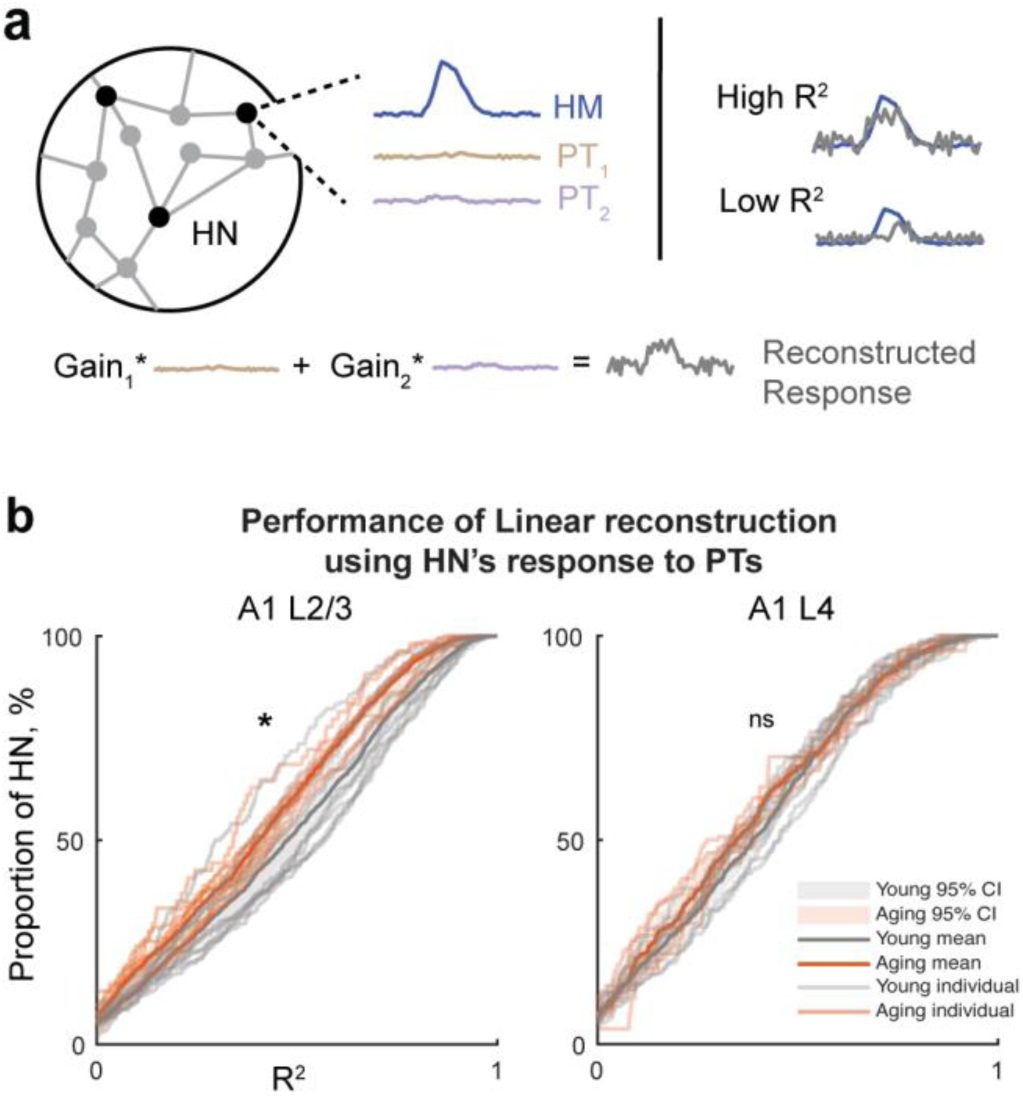
Linearity of spectral integration by HNs in A1 L2/3 and L4. **a.** Schematic showing the linear reconstruction of HN’s response to the harmonic stack by the HN’s response to the pure-tones (PTs). An example of high R^2^ (0.65) shows high overlap of the reconstructed response to the original response. An example of low R^2^ (0.15) shows poor overlap of the reconstructed response to the original response. **b.** Cumulative distribution curves that show the distribution of R^2^ of all HNs for individual animals (thin lines) and the average curves of young (thick red line) and aging (thick grey line) cohorts in A1 L2/3 (left) and L4 (right). Permutation tests (5000 iterations) on animal IDs were used to assess significance. *p*L2/3 = 0.026, *p*L4 = 0.33. * indicates *p* 0 0.05. ns, no significance.

**Extended Figure 3:**
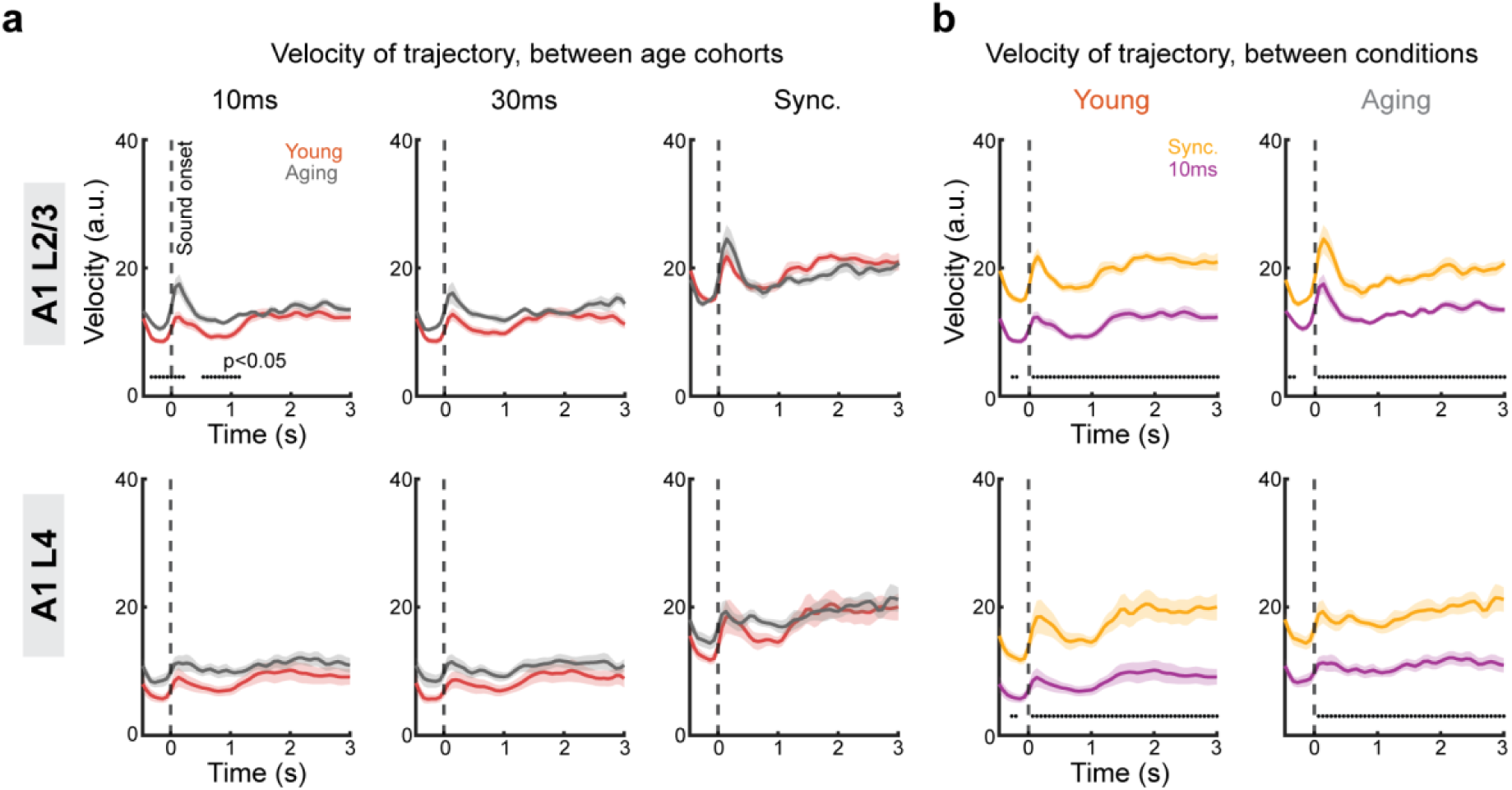
Neural trajectory velocity in ACtx across age groups and sound conditions. **a.** Mean trajectory velocity (±SEM as shaded area) in the neural state space for young (red) and aging (gray) animals, shown separately for each stimulus condition (10ms shift, 30ms shift, and synchronous harmonics) in ACtx L2/3 (top) and L4 (bottom). Each velocity value was calculated as the difference of distance at current time frame from the previous time frame. Dashed vertical line indicates stimulus onset. Significant differences from two samples t-tests between age groups are marked as black dots where *p*00.05 after Benjamini-Hochberg corrections for multiple comparisons at each time frame. **b.** Mean trajectory velocity across conditions (Sync, yellow; 10ms shift, purple) compared between young (left) and aging (right) animals for L2/3 (top) and L4 (bottom). Shaded regions indicate ±SEM. Significant differences from two samples t-tests between age groups are marked as black dots where *p*00.05 after Benjamini-Hochberg corrections for multiple comparisons at each time frame.

**Extended Figure 4:**
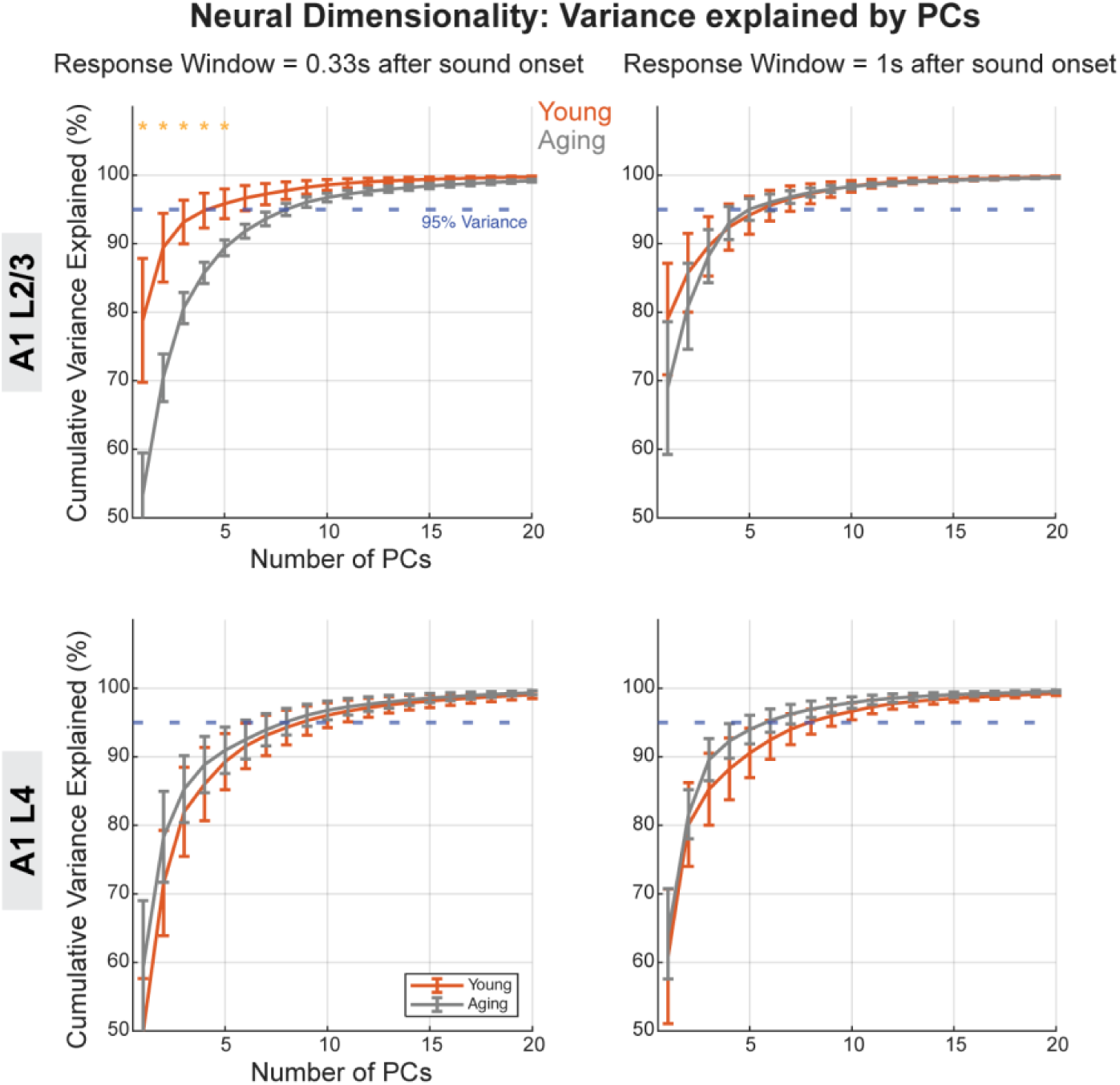
Variance explained by principal components (PC) using short and long response window in L2/3 and L4. Top: average variance explained by varied numbers of PCs computed based on 0.33s response window after sound onset (left) and 1s response window after sound onset (right) in A1 L2/3. Two-sample t-tests were used at each number of PCs. Yellow * indicates significant difference (*p* 0 0.05 after Benjamini-Hochberg correction) between young and aging cohorts on the cumulative variance. Error bars indicate SEM. Bottom: Same as Top, but in A1 L4.

**Extended Figure 5:**
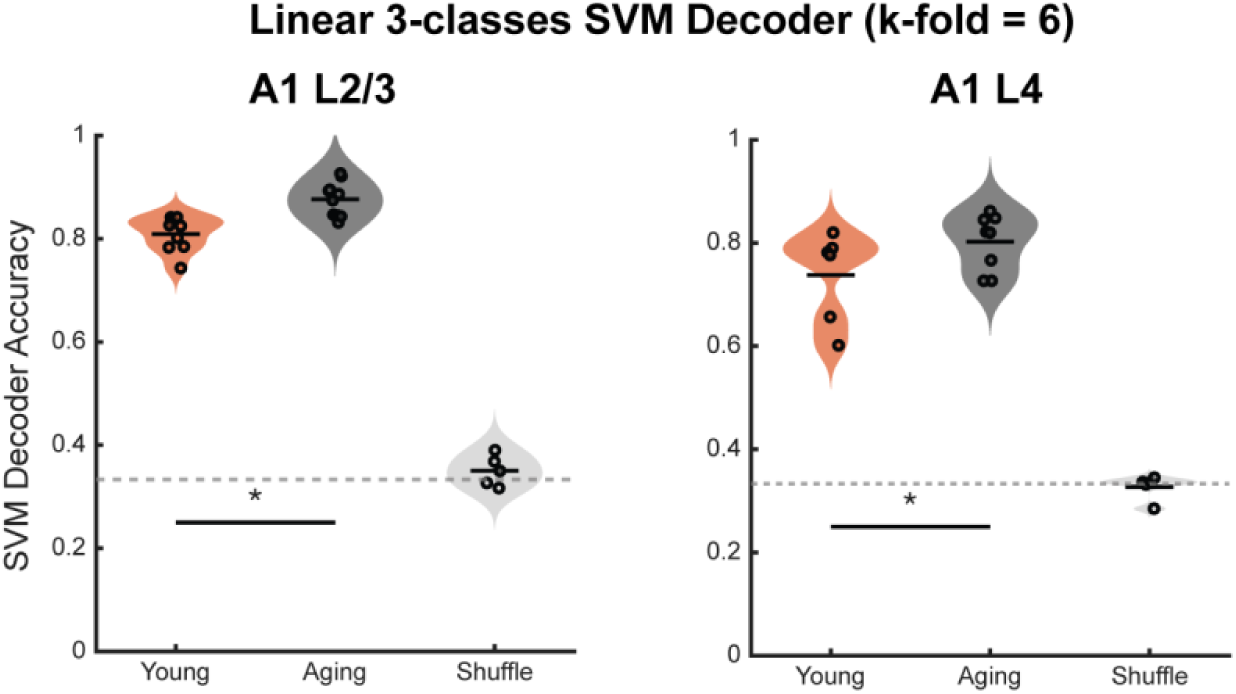
Accuracies of a linear 3-classes SVM decoder. Accuracies of using a linear 3-classes SVM decoder with k-fold=6 validation to classify trial types in L2/3 (left) and L4 (right), respectively, based on response described in Fig. 2f. Data input was identical to those used in Fig. 2h except the type of decoder. Dashed horizontal line indicates chance-level performance based on shuffle controls. Mann-Whitney U test was used. YL2/3: 80.93% ±1.12%, AL2/3: 87.65% ±1.07%, *p*L2/3, A vs Y = 1.30e-4. YL4: 73.78% ±3.56%, AL4: 80.21% ±1.93%, *p*L4, A vs Y = 0.15.

## Notes

**Conflicts of interest:** No conflicts.

### Competing Interest Statement

The authors have declared no competing interest.

